# Neutrophil extracellular traps block endogenous and intravenous thrombolysis-induced fibrinolysis in large vessel occlusion acute ischemic stroke

**DOI:** 10.1101/2025.03.28.646062

**Authors:** Jean-Philippe Desilles, Mialitiana Solo Nomenjanahary, Astride Perrot, Lucas Di Meglio, Fatima Zemali, Sara Zalghout, Stéphane Loyau, Julien Labreuche, Marie-Charlotte Bourrienne, Dorothée Faille, François Delvoye, Véronique Ollivier, Sébastien Dupont, Jasmina Rogozarski, Nahida Brikci-Nigassa, Nadine Ajzenberg, Mikael Mazighi, Benoît Ho-Tin-Noé, the compoCLOT study group

**Affiliations:** Université Paris Cité, INSERM, Optimisation Thérapeutique en Neuropharmacologie OTEN U1144, 75006 Paris, France; Biological Resource Center and Department of Interventional Neuroradiology, Rothschild Foundation hospital, Paris, France; Université Paris Cité, Inserm, UMRS-1148, Laboratory for Vascular Translational Science, 75018 Paris, France; CHU Lille, Department of Biostatistics, F59000-Lille, France; FHU NeuroVasc, Paris, France; Department of Neurology, GHU APHP NORD, Hôpital Lariboisière, 2 rue Ambroise Paré, 75010, Paris, France; Stroke-Link F-CRIN research network; Institut Universitaire de France

**Keywords:** intravenous thrombolysis, thrombolysis resistance, neutrophil extracellular traps, ischemic stroke, DNase 1, fibrinolysis

## Abstract

**Background:** Intravenous thrombolysis (IVT) failure in acute ischemic stroke (AIS) due to large vessel occlusion (LVO) is frequent but its causes remain elusive. Several non-exclusive mechanisms have been proposed to explain IVT failure, including failed delivery of tPA and inhibition of its activity. We investigated whether biologically relevant intrathrombus concentrations of t-PA were achieved in failed IVT in patients with LVO AIS, and whether neutrophil extracellular traps (NETs) contributed to IVT failure.

**Methods:** In this cohort study, a total of 205 thrombi from AIS patients with LVO were analyzed. 83 of these thrombi were compared for tPA content and 53 for their susceptibility to *ex vivo* thrombolysis according to IVT status. An additional subset of 69 AIS thrombi was used to decipher if and how NETs interfere with intrathrombus fibrinolysis.

**Results:** AIS thrombi from IVT patients contained more tPA than those from no-IVT patients (0.209 vs 0.093 µg/mg of thrombus, p<0.0001). Plasminogen and tPA in AIS thrombi were found in association with fibrin and NETs. The ability of NETs to bind tPA and plasminogen, titrating them away from fibrin, was confirmed in a microfluidic model of thrombosis. While *ex vivo* addition of plasminogen did not cause lysis of either no-IVT or IVT thrombi, combining plasminogen with DNase 1 helped translate the increased tPA content of IVT thrombi into increased thrombolysis. We further show that DNase 1 enables tPA- and plasmin-mediated thrombolysis by eliminating fibrinolysis inhibitors from AIS thrombi.

**Conclusions:** These results indicate that intrathrombus tPA concentrations reached in failed IVT bear a therapeutic potential that is however impaired by NETs, which favor intrathrombus retention of fibrinolysis inhibitors and compete with fibrin for tPA and plasminogen binding. Our results stress the interest of DNase 1 to enhance the efficacy of current IVT tPA regimens.

**Clinical Perspective:** *What is new?:*

- Intravenous thrombolysis increases thrombus tPA content even when it fails to cause arterial recanalization in acute ischemic stroke
- The fibrinolytic activity of intravenously-administered tPA is blocked by neutrophil extracellular traps in acute ischemic stroke thrombi
- Neutrophil extracellular traps participate in thrombolysis resistance by retaining fibrinolysis inhibitors and titrating tPA and plasminogen away from fibrin in acute ischemic stroke thrombi
- DNase 1 can convert increased tPA content into increased fibrinolysis by eliminating NETs-associated fibrinolysis inhibitors in acute ischemic stroke thrombi

*What are the clinical implications?:*

- Despite therapeutic failure, biologically significant intrathrombus tPA concentrations are achieved following intravenous thrombolysis at current tPA regimens
- Sequential administration of DNase 1 prior to intravenous thrombolysis could clear the way for tPA and potentiate its fibrinolytic activity for improved arterial recanalization efficacy

## Introduction

The recanalization efficacy of intravenous thrombolysis (IVT) with recombinant tissue-type plasminogen activator (tPA) in acute ischemic stroke (AIS) patients is particularly limited in case of large vessel occlusion (LVO)^1^. However, IVT-induced recanalization in AIS patients before endovascular therapy (EVT) is significantly associated with better clinical outcomes^2^. The proximal location of the AIS thrombus is one of the main predictors of poor IVT efficacy. In addition to thrombus localization, imaging studies have indicated that thrombus size also influences IVT efficacy, with larger thrombi being more resistant to IVT^3,4^. Collateral status may be an important factor as well^5,6^, presumably by modulating tPA delivery into the thrombus. Nonetheless, to date, there are no data on intrathrombus tPA content following failed IVT. Such data would be helpful to determine whether IVT failure is associated with failed delivery of tPA to AIS thrombi.

Analyses of EVT-recovered thrombi have indicated that apart occlusion site, thrombus length, and collateral status, thrombus structure and composition may also modulate IVT efficacy^7^. In particular, platelets and neutrophil extracellular traps (NETs) have been identified as potential inhibitors of IVT^8–14^. Interestingly, multiple groups have shown that the thrombolysis of AIS thrombi by tPA added *ex vivo* can be increased by the concomitant addition of recombinant DNase 1 to degrade intrathrombus NETs^8,9,11,12,14^. These data have indicated that the degradation of nonfibrin fibrillar thrombus components like NETs and von Willebrand factor could help bypass resistance to tPA but it also raised the question of a possible inhibition of the fibrinolytic activity of tPA by these components. Whether NETs block the action of tPA in AIS due to LVO, and whether DNase 1 could improve the thrombolytic effect of endogenous or IVT-administered tPA remains unknown.

In the present study, we investigated whether biologically relevant intrathrombus concentrations of t-PA were achieved following IVT in patients with LVO-related AIS, and if and how NETs contributed to IVT failure.

## Methods

### Standard protocol approvals, registrations, and patient consents

Thrombi were collected at the end of EVT. The EVT procedure was chosen at the interventionalist’s discretion using a stent-retriever and/or a contact aspiration technique. Patient data were collected prospectively using a standardized questionnaire (Endovascular Treatment in Ischemic Stroke-ETIS-registry NCT03776877). All patients and healthy blood donors were provided with a written explanation of the study. The patients or their representatives were given the opportunity to refuse participation. The local Ethics Committee approved this research protocol (CPP Nord Ouest II, ID-RCB number: 2017-A01039-44).

### Acute ischemic stroke thrombi collection and processing

A total of 205 thrombi recovered by EVT between August 2016 and July 2024 were included in this study. Of those, 83 thrombi (43 no-IVT, 40 IVT) were used for preparation of thrombus homogenates, 53 (24 no-IVT, 29 IVT) were for *ex vivo* thrombolysis assays according to IVT status, 10 were fixed in paraformaldehyde upon collection and used for immunohistological analysis, and 59 were used for sequential *ex vivo* thrombolysis assays. Each thrombus was randomly allocated to one of these investigation methods.

### Immunostaining

Within 60 min of EVT completion, thrombi were fixed in 3.7% paraformaldehyde for 48 h before being embedded longitudinally in paraffin and sectioned at 5 μm. Immunostainings were performed as described previously using primary antibodies to fibrinogen (A0080, Dako), myeloperoxidase (A0398, Dako), tPA (HTPA2A153, Molecular Innovation), plasminogen (80 µg/mL, PA5-14196, Invitrogen), plasminogen activator inhibitor-1 (PAI-1) (MA-33B8, Merck), alpha-2-antiplasmin (ab150414, abcam). DNA was identified using Hoechst 33342 (H3570, Life Technologies) and red blood cells (RBCs) were identified by their autofluorescence (λex 440/9/λem>570 nm).

### *Ex vivo* thrombolysis assays

Thrombi were cut in two equal parts and treated with or without DNase 1 (100 μg/ml, Dornase Alfa, Pulmozyme; Roche, Basel, Switzerland) at 37°C under agitation (500 rpm, Thermomixer) in D-PBS supplemented with plasminogen (100 µg/mL 1 μM, Technoclone; Vienna, Austria). A 25-μl aliquot of lysis supernatant was collected 10 min after eliciting thrombolysis for measurement of D-dimers. A subset of thrombi was incubated with DNase 1 ± aprotinin (Trasylol, 2 µM), without addition of plasminogen.

For sequential thrombolysis, AIS thrombi were cut in half, thrombus fragments were then first incubated for 2 hours with either 100 µg/mL DNase 1 (pulmozyme) or corresponding volume of vehicle (1 mM calcium chloride in saline) prepared in D-PBS, rinsed twice with after collecting the supernatants, and thrombolysis was then elicited by addition of tPA (1 μg/ml Alteplase, Actilyse; Boehringer Ingelheim, Ingelheim am Rhein, Germany) and plasminogen (100 µg/mL), plasmin (200 nM), or plasminogen/streptokinase. For preparation of plasminogen/streptokinase, plasminogen was diluted at 20 µg/mL in D-PBS containing streptokinase in excess (5000 IU/mL, STAGO). All incubation volumes were adjusted to the thrombus weight at a v/w ratio of 40 ml/g.

### Measurement of tPA, plasminogen, glycoprotein VI, DNA, hemoglobin, D-dimers in homogenates from no-IVT or IVT thrombi

Homogenates from no-IVT and IVT thrombi were prepared as described previously^15^. Plasminogen and tPA content were quantified by ELISA (Technozym Glu-plasminogen TC12040, Technoclone and Asserachrom tPA, 00948, Stago) following the manufacturers’ instruction. DNA was quantified using Quant iT Picogreen dsDNA Assay kit (Molecular Probes, Life Technologies). Thrombus platelet content was estimated using a mesoscale-based immunoassay for glycoprotein VI (GPVI), and RBC content by measurement of heme concentration using a formic acid-based colorimetric assay, as described previously^15,16^. D-dimers were measured by quantitative latex immunoassay (HemosIL D-dimer HS500, Werfen) on an ACL TOP 700 analyzer (Werfen).

### Measurement of fibrinolysis inhibitors in supernatants from DNase 1-treated ischemic stroke thrombi

Alpha-2-antiplasmin (α2AP, DuoSet Human Serpin2/alpha-2-antiplasmin, DY1470, R&D Systems), PAI-1 (Asserachrom 00949, Stago), and neutrophil elastase (Hycult Biotech HK319) were quantified in supernatants from AIS thrombi using commercial kits and following the manufacturers’ instructions. Protease nexin-1 was measured as described previously^17^.

### Real-time video microscopic analysis of tPA and plasminogen interactions with a growing thrombus

Citrated whole blood from healthy donors was recalcified with calcium chloride to reach 0.35 ±0.05 mM of ionised calcium, spiked with glucose to reach a glycemia of 3g/L, supplemented with Hoechst 33342 and Alexa Fluor-conjugated antibodies to fibrin (1 µg/mL, clone 59D8, AntibodySystem, ref RHB99201) and plasminogen (2.5 µg/mL, clone HL2071, Invitrogen, ref MA5-47278), before being perfused at a flow rate of 12 µL/min (corresponding to a shear stress of 13.5 dyne/cm^2^ and a wall shear rate of 300 s^-1^), through Vena8 Fluoro+ biochip channels (Cellix) coated with fibrin monomers prepared as described with adaptations^18^. Thrombus and NETs formation were monitored in real-time by fluorescence microscopy (DMi8 microscope, Leica), using a sCMOS camera operated by LasX software (Leica). Once a NETs-rich thrombus had formed, the whole blood perfusate was supplemented with Alexa Fluor-conjugated tPA (10 µg/mL, Alteplase, Actilyse; Boehringer Ingelheim).

### Western blotting

Proteins from thrombus homogenates loaded in non-reducing and reducing Laemmli buffer were resolved by polyacrylamide gel electrophoresis (8% and 4-12%) and transferred to polyvinylidene difluoride membranes using the ThermoFisher iBlot 3 Western Blot Transfer System. The membranes were probed with antibodies to α2AP (ab150414, abcam), plasminogen (PA5-14196, Invitrogen), tPA (NBP2-20648, Novus Biologicals), and histone H1 (SC-67324, Santa Cruz), revealed with Dylight 800- and Dylight 680-conjugated secondary antibodies using a LI-COR Odyssey® imaging system.

### Statistical analysis

Categorical variables were expressed as frequencies and percentages. Quantitative variables were expressed as mean (standard deviation) or median (25th to 75th percentiles) in case of non-normal distribution. Normality of distributions was assessed using histograms and the Shapiro-Wilk test. Patients Baseline characteristics and thrombus cell marker contents were described according to intravenous thrombolysis use prior to mechanical thrombectomy; the magnitude of the between-group differences was assessed by calculating the absolute standardized differences (ASD); an ASD>20% in was interpreted as a meaningful difference^19^. Comparisons in thrombus fibrinolytic enzymes and markers between IVT and non-IVT treated patients were done using Mann-Whitney U tests. Among IVT treated patients, association of DDimers and tPA contents with several factors were assessed using Mann-Whitney test for binary factors or by calculating the Spearman’s rank correlation coefficients for quantitative factors. The treatment effects in ex vivo thrombolysis assays were assessed using the Wilcoxon signed-rank test for paired samples or using Mann-Whitney U test for independent samples. Statistical testing was conducted at the two-tailed α-level of 0.05 without correction for multiple comparisons due to the exploratory nature of this study. Data were analyzed using the SAS software version 9·4 (SAS Institute, Cary, NC) or using PrismGraph 4.0 software (GraphPad Software, San Diego, CA).

## Results

### Intravenous thrombolysis increases tPA and d-dimer content in AIS thrombi despite recanalization failure

In this cohort study, a total of 83 thrombi from AIS patients with LVO having received (n=40, IVT) or not (n=43, no-IVT) IVT prior to EVT were analyzed and compared for tPA, D-dimer, and plasminogen content, and for cell composition, as assessed by quantitative measurement of glycoprotein VI (GPVI), DNA, and heme content, used as markers of platelets, leukocytes, and red blood cells, respectively. There was no significant difference in baseline characteristics between IVT and no-IVT patients, or in their GPVI, DNA, heme, and plasminogen thrombus content (Table 1 and Figure 1). Thrombi from IVT patients had significantly higher levels of tPA and D-Dimer compared to thrombi from no-IVT patients (Supplemental Table 1, Figure 1A and B). These results indicate that IVT does increase thrombus tPA content and initiates fibrinolysis despite failure of thrombolysis-induced recanalization. Intrathrombus tPA content in IVT patients was negatively correlated with the delay between IVT and mechanical thrombectomy-mediated thrombus recovery (Supplemental Table 1). Western blot analysis of tPA in IS thrombus homogenates showed that, in both no-IVT and IVT thrombi, tPA was mainly present as high-molecular-weight complexes, one of which whose size of approximately 110 kDa was compatible with that of PAI-1/tPA complexes (Figure 1C). The highest molecular weight complex was also positive for histone H1, indicating that tPA formed complexes involving histones as well (Supplemental Figure 1). All complexes were dissociated upon reducing conditions, yielding single chain tPA (Figure 1C).

**Figure 1.**
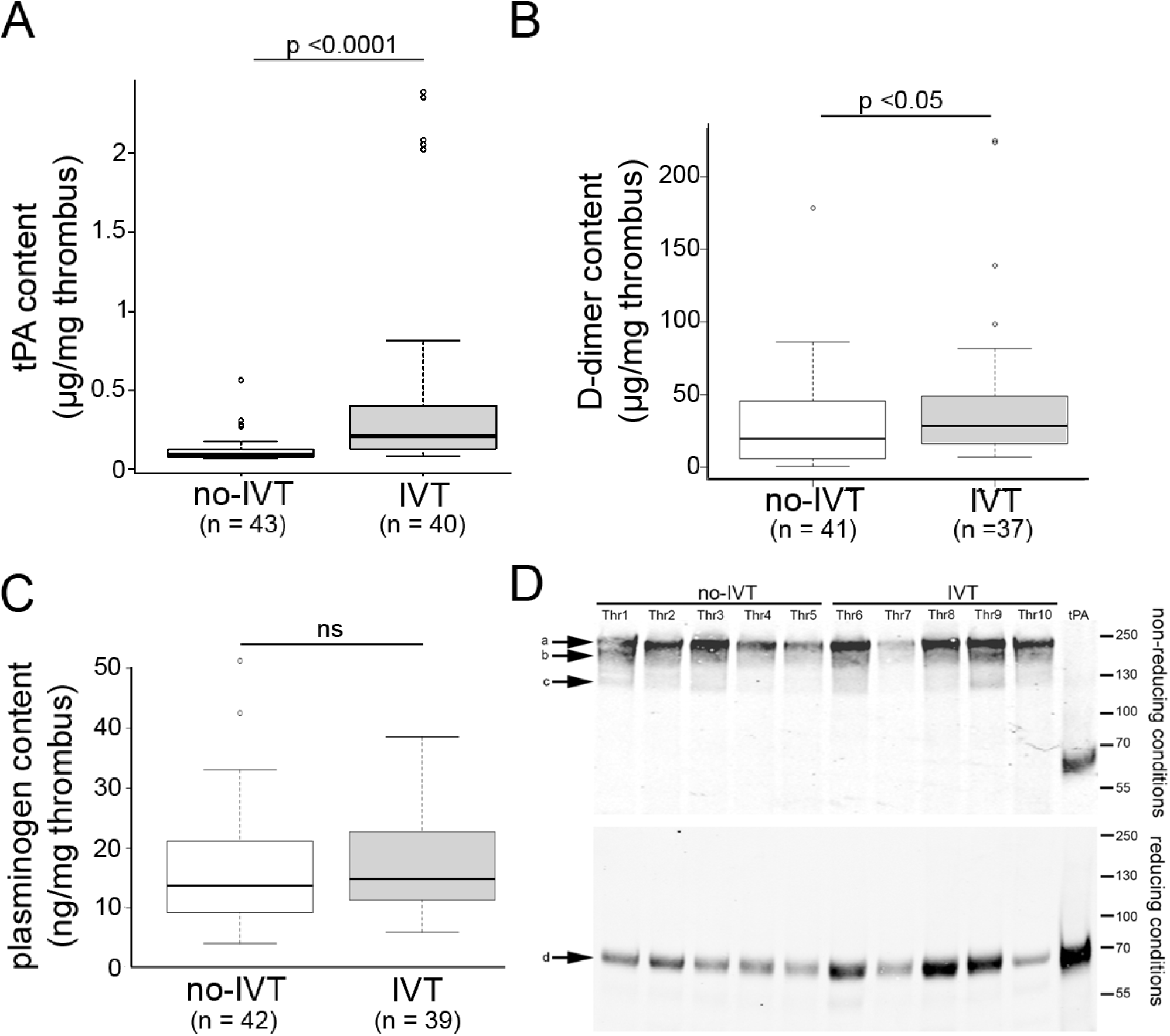
Intravenous thrombolysis increases tissue-type plasminogen activator content in acute ischemic stroke thrombi despite therapeutic failure. A-C. Comparison of tissue-type plasminogen activator (tPA) (A), D-dimer (B), and plasminogen (C) content in thrombi from patients with acute ischemic stroke due to large vessel occlusion having received intravenous thrombolysis (IVT) or not (no-IVT) before endovascular thrombectomy. D. Western blot analysis of tPA in homogenates of 5 no-IVT (Thr1 to 5) and 5 IVT (Thr6 to 10) thrombi in non-reducing (upper panel) and reducing (lower panel) conditions. Note that in both no-IVT and IVT thrombi, tPA is mainly found under the form of 3 high-molecular-weight complexes (indicated by arrows and labeled a to c in the upper panel), from which it could be dissociated as free single chain tPA under reducing conditions (arrow and labeled d in the lower panel).

**Table 1.** Baseline characteristics and thrombus cell marker content according to intravenous thrombolysis use prior to mechanical thrombectomy. Values expressed as no/total no. (%) unless otherwise indicated. *1 missing data (0 in IV-treated patients); ^†^6 missing data (3 in IV-treated patients); ^‡^2 missing data (1 in each group); ^§^5 missing data (3 in IV-treated patients). Abbreviations: ASD=absolute standardized difference; GPVI = glycoprotein VI; ICA = internal carotid artery; IQR = interquartile range; MCA = middle cerebral artery; NIHSS = National Institutes of Health Stroke Scale; tPA= tissue plasminogen activator; TIA = transient ischemic attack; mRS = modified Rankin scale, SD = standard deviation.

| Characteristics | Intravenous thrombolysis |  | ASD |
| --- | --- | --- | --- |
|  | No (n=43) | Yes (n=40) |  |
| Demographics |  |  |  |
| Age, years, mean (SD) | 72.3 (16.7) | 66.4 (15.3) | 37.0 |
| Men | 21/43 (48.8) | 17/40 (42.5) | 12.8 |
| Medical history |  |  |  |
| Hypertension | 29/43 (67.4) | 20/39 (51.3) | 33.4 |
| Diabetes | 11/43 (25.6) | 5/39 (12.8) | 32.8 |
| Hypercholesterolemia | 13/42 (31.0) | 9/39 (23.1) | 17.8 |
| Current smoking | 9/40 (22.5) | 5/39 (12.8) | 25.6 |
| Coronary artery disease | 6/43 (14.0) | 4/39 (10.3) | 11.4 |
| Previous stroke or TIA | 9/43 (20.9) | 1/38 (2.6) | 59.2 |
| Previous antithrombotic medications | 19/43 (46.3) | 10/40 (25.0) | 45.7 |
| Current stroke event |  |  |  |
| NIHSS score, median (IQR) | 18 (16 to 22) | 19 (13 to 22) | 8.4 |
| Pre-stroke mRS≥1 | 10/42 (23.8) | 5/40 (12.5) | 29.7 |
| Intracranial ICA occlusion | 19/42 (45.2) | 15/40 (37.5) | 34.4 |
| Cardioembolic etiology | 26/43 (60.5) | 20/40 (50.0) | 21.2 |
| Onset to groin puncture time, min<sup>*</sup> | 251 (181 to 335) | 264 (207 to 292) | 7.5 |
| Thrombus weight, median (IQR) | 63 (43 to 108) | 57 (38 to 82) | 5.3 |
| Thrombus cell marker content |  |  |  |
| DNA, µg/mg thrombus, median (IQR) | 28 (12 to 94) | 41 (17 to 106) | 15.8 |
| GPVI, µg/mg thrombus, median (IQR)<sup>†</sup> | 0.108 (0.090 to 0.118) | 0.103 (0.089 to 0.110) | 33.1 |
| HEME, µg/mg thrombus, median (IQR) | 210 (155 to 301) | 232 (159 to 254) | 1.8 |
| Thrombus fibrinolytic enzymes and markers |  |  |  |
| tPA, µg/mg thrombus, median (IQR) | 0.093 (0.075 to 0.129) | 0.209 (0.128 to 0.404) | 145.0<sup>*</sup> |
| Plasminogen, ng/mg thrombus, median (IQR)<sup>‡</sup> | 22.4 (13.7 to 34.4) | 24.4 (13.9 to 32.8) | 0.4 |
| D-dimers, µg/mg thrombus, median (IQR)<sup>§</sup> | 19200 (5546 to 45270) | 28320 (15780 to 48616) | 46.2<sup>*</sup> |

Immunostaining for tPA in IS thrombi confirmed the presence of tPA in IS thrombi irrespective of IVT status. As expected, tPA was found in association with fibrin, both at the thrombus periphery and in the core, but also in association with NETs (Figure 2). Likewise, plasminogen was found in association with fibrin, NETs, and neutrophil surface (Figure 3). Notably, the ability of NETs to bind plasminogen and tPA was confirmed *in vitro* in a microfluidic model of thrombosis. Real-time video microscopic analysis of plasminogen and tPA interactions with a growing thrombus showed that, as indicated by immunostaining of IS thrombi (Figure 2), plasminogen and tPA bound to fibrin but also accumulated in NETs-riched areas (Supplemental Video 1 and 2).

**Figure 2.**
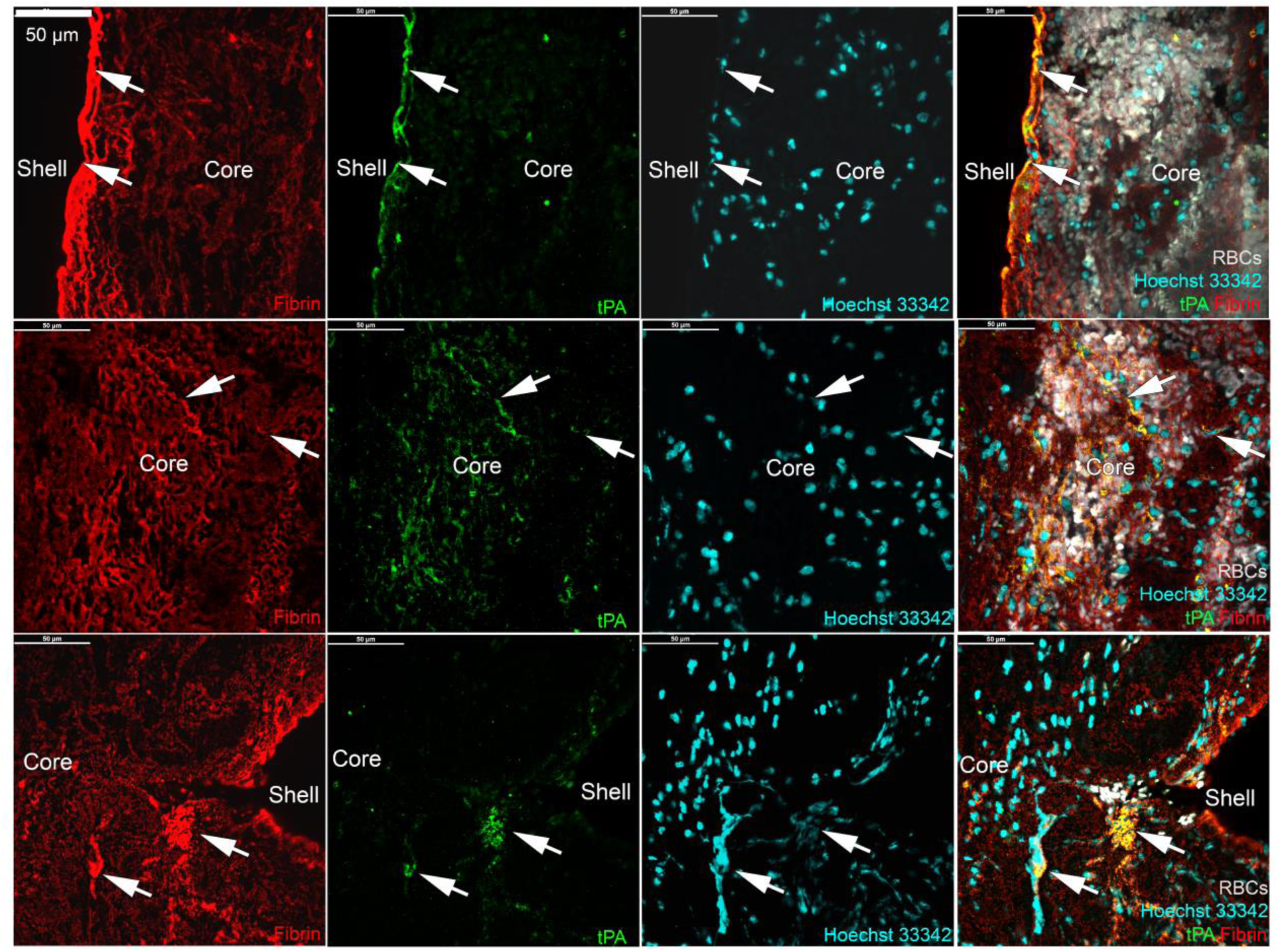
Localization of tissue-type plasminogen activator in acute ischemic stroke thrombi. **A.** Representative images of immunofluorescent localization of tissue-type plasminogen activator (tPA, green) in thrombi from IS patients. Top panels: accumulation of tPA in association with compact fibrin (red) in the thrombus outer shell (arrows). Middle panels: tPA bound to fibrin fibers in a red blood cell-rich area of the thrombus core and with compact fibrin (arrows). Lower panels: the white arrows indicate tPA associated with clusters of dense fibrin entangled with extracellular DNA (cyan) close to the thrombus surface. The images shown were acquired in a thrombus from a patient having received intravenous thrombolysis prior to EVT and are representative of 10 different thrombi.

**Figure 3.**
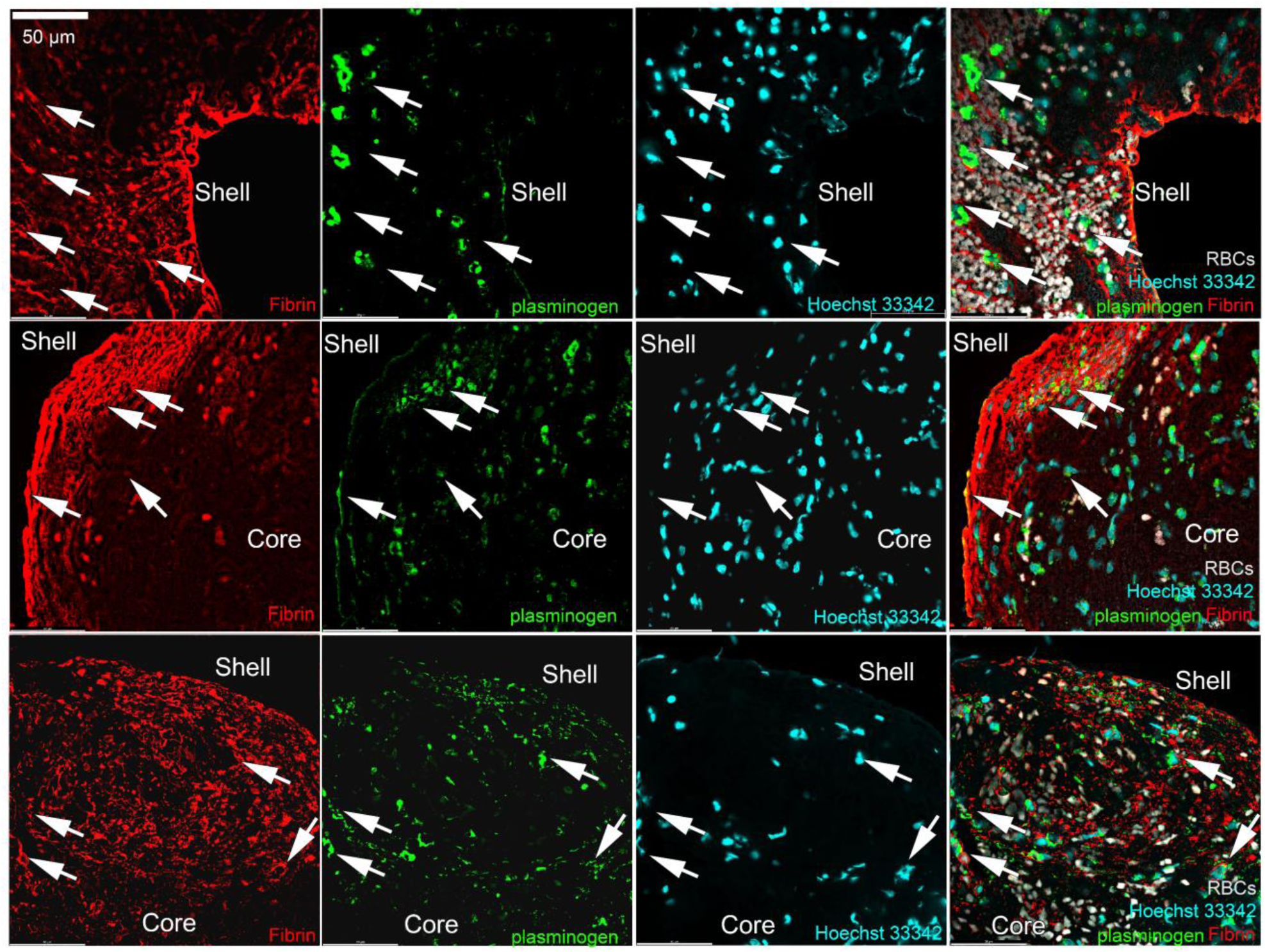
Localization of plasminogen in acute ischemic stroke thrombi. Representative images of immunofluorescent localization of plasminogen (green) in various thrombi from acute ischemic stroke patients showing the association of plasminogen with dense fibrin (red), nucleated cells and NETs (cyan, arrows).

Western blot analysis of intrathrombus plasmin(ogen) content and structure did not show differences according to IVT status. In non-reducing conditions, plasmin(ogen) was detected under 3 different forms in both IVT and no-IVT thrombi, a high molecular band of approximately 190 kDa, a 90 kDa band corresponding to the expected molecular weight of plasmin(ogen), and a lower molecular weight band, whose size (∼ 35-40 kDa) matched that of previously described neutrophil elastase-mediated plasminogen fragments^20–22^ (Supplemental Figure 2). In reducing conditions, the high molecular band was lost and the plasminogen band faded in association with the appearance of the heavy and light chains of plasmin, indicating that a fraction of intrathrombus plasminogen had been converted to plasmin in both IVT and no-IVT patients (Supplemental Figure 2).

### *Ex vivo* addition of DNase 1 enables endogenous and intravenous thrombolysis-related fibrinolysis in ischemic thrombi

Because tPA induces thrombolysis by activating plasminogen into plasmin, we investigated whether recanalization failure despite increased tPA content following IVT was due to insufficient plasminogen. *Ex vivo* addition of plasminogen at 1 µM did not cause any reduction in the weight of either no-IVT or IVT AIS thrombi over a 60-min-long incubation period (Figure 4A-C), indicating that factors other than plasminogen concentration contributed to IVT failure. We have shown previously that the ability of DNase 1 to enhance the lysis of AIS thrombi by tPA added *ex vivo^9,^*^11,12^ is associated with increased fibrinolysis^23^. We thus tested whether DNase 1 (dornase-alfa, pulmozyme, Roche) could help translate the increased tPA content of IVT thrombi into significant thrombolysis. Combined addition of DNase 1 and plasminogen caused a significant decrease in the weight of both no-IVT- and IVT AIS thrombi over a 60-min-long lysis period (Figure 4A-C), which was more pronounced in IVT AIS thrombi (Figure 4B-C). The increased lysis in the presence of DNase 1 was associated with an increase in fibrinolysis, as indicated by increased D-Dimer release (Supplemental Figure 3). When comparing the impact of DNase1 and plasminogen according to thrombolytic drug used for IVT, there was a non-significant trend for increased lysis with tenecteplase vs alteplase (Figure 4D).

**Figure 4.**
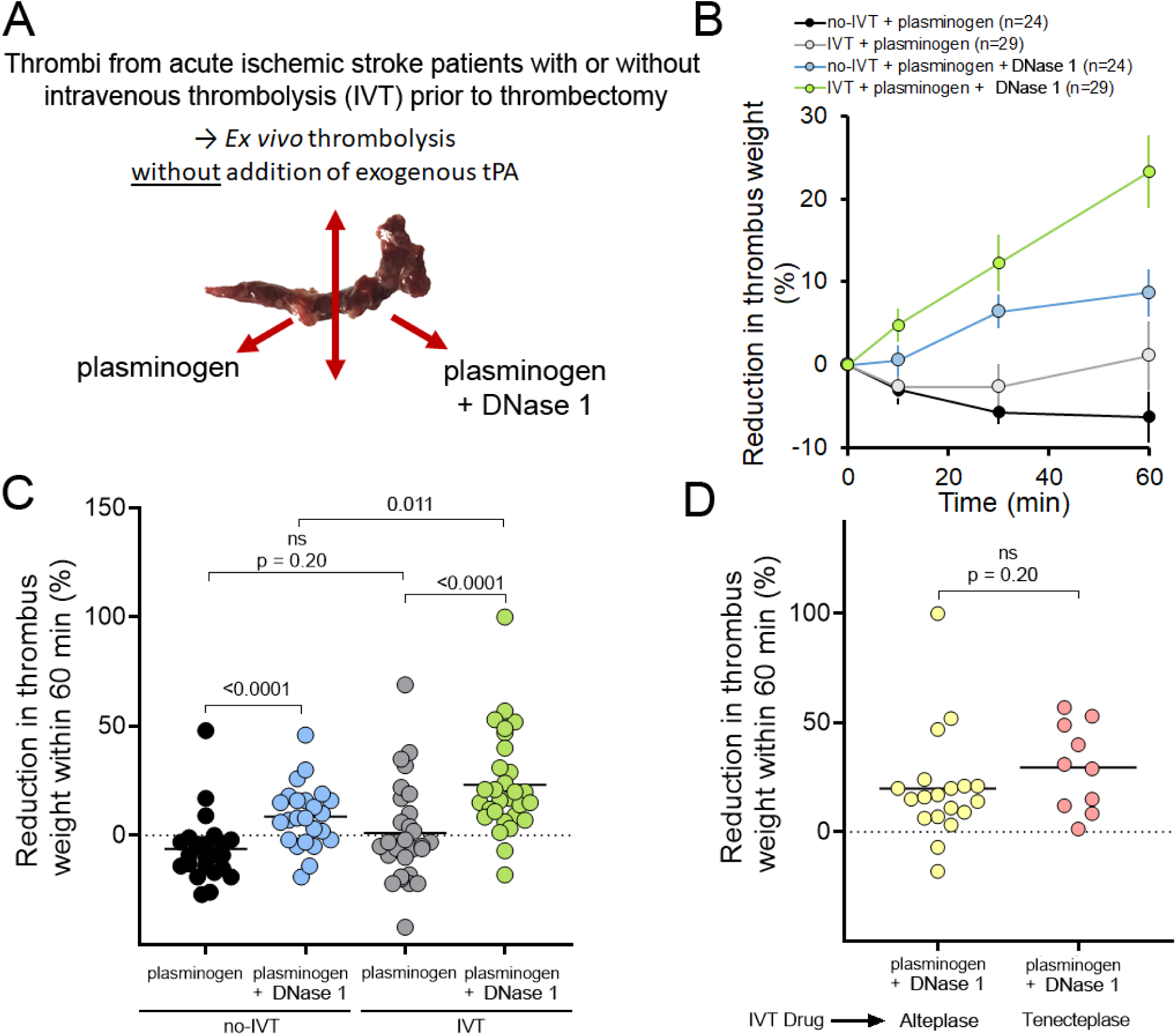
DNase 1 potentiates the thrombolytic activity of tissue-type plasminogen activator delivered to acute ischemic stroke thrombi by intravenous thrombolysis. **A.** Schematic representation of the *ex vivo* thrombolysis assay performed with thrombi from stroke patients that had or had not received intravenous thrombolysis with tissue-type plasminogen activator (tPA) prior to mechanical thrombectomy. **B.** Mean thrombus weight evolution over 1 hour of *ex vivo* thrombolysis initiated by the addition of plasminogen (100 µg/mL), in the presence or absence of DNase 1 (pulmozyme, 100 µg/mL), between thrombi from patients having received (IVT) or not (no-IVT) intravenous thrombolysis prior to thrombectomy. Error bars indicate standard error of the mean. **C.** Comparison of the reduction in thrombus weight after 60 min of *ex vivo* incubation of acute ischemic thrombi (AIS) with plasminogen alone or combined with DNase 1 between thrombi recovered from patients having received tPA (IVT) or not (no-IVT) prior to thrombectomy. Results are expressed as a percentage relative to the initial thrombus weight. **D.** Comparison of the reduction in thrombus weight after 60 min of *ex vivo* incubation of AIS thrombi with plasminogen and DNase 1 according to the type of recombinant tPA used for IVT. Results are expressed as a percentage relative to the initial thrombus weight. Each dot represents a different thrombus.

Another subset of IS thrombi was used to determine if DNase 1 had thrombolytic effects irrespective of fibrinolysis. For these experiments, 14 IS thrombi were split in half, one half was treated with DNase 1 without addition of exogenous plasminogen; the other half received the same treatment supplemented with aprotinin in order to block fibrinolysis that could arise from plasminogen and tPA present in AIS thrombus. No reduction in thrombus weight occurred in either condition after 60 min of treatment. However, after 120 min of treatment, DNase 1 alone had caused a mean reduction in thrombus weight of 7.4 % (Supplemental Figure 4). Yet, this mean reduction in thrombus weight was prevented in the presence of the fibrinolysis inhibitor aprotinin (Supplemental Figure 4). When considering individual AIS thrombus response to DNase 1, only 2 out of the 14 thrombus fragments showed substantial weight loss despite the presence of aprotinin, thus indicating that DNase 1 causes thrombolysis irrespective of fibrinolysis only in a minority of AIS thrombi. Taken together, these results show that although DNase 1 has little direct thrombolytic effect, it can unlock the fibrinolytic potential of both endogenous and IVT-delivered tPA in AIS thrombi.

### DNase 1 enhances tPA-mediated thrombolysis by waiving NETs-dependent intrathrombus retention of fibrinolysis inhibitors and promoting alpha-2-antiplasmin degradation

Using a subset of IS thrombi, we then assessed whether the pro-fibrinolytic effect of DNase 1 was maintained when tPA was added after DNase 1 treatment and elimination by washing of potential soluble factors released during DNase 1 treatment (Figure 5A). The *ex vivo* lysis of IS thrombi by tPA and plasminogen was markedly increased following a 2-hour-long pretreatment period with DNase 1 as compared to preincubation with vehicle (Figure 5B-C). Together, these results indicate that enhancement of tPA-mediated thrombolysis by DNase 1 does not rely on the release of profibrinolytic soluble factors and instead suggest that DNase 1 might help eliminate antifibrinolytic factors.

**Figure 5.**
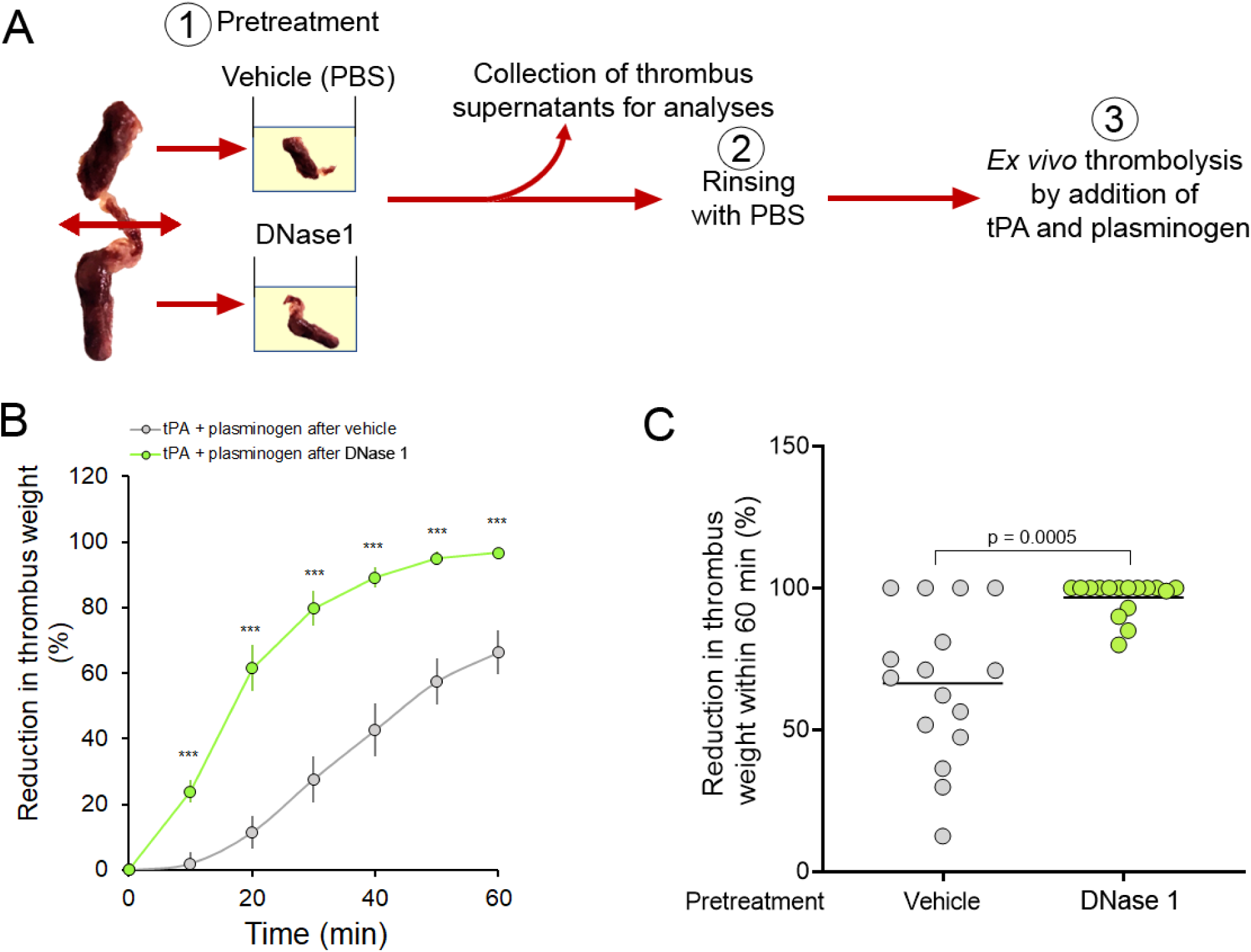
DNase 1 pretreatment facilitates subsequent thrombolysis of acute ischemic stroke thrombi by tissue-type plasminogen activator. **A.** Schematic representation of sequential *ex vivo* thrombolysis in which acute ischemic stroke (AIS) thrombi were cut in half and first incubated with DNase 1 alone (pulmozyme, 100 µg/mL), before being subsequently treated with tissue-type plasminogen activator (tPA, alteplase, 1 µg/mL) and plasminogen (100 µg/mL or 1 µM). **B.** Comparison of mean thrombus weight evolution in response to tPA and plasminogen added following a pretreatment with either DNase 1 or vehicle. n = 16 thrombi, *** p < 0.0005 for the comparison between the two groups at each time point. **C.** Paired comparison of thrombus weight after 60 min of *ex vivo* thrombolysis triggered by addition of tPA and plasminogen after a pretreatment of AIS thrombi with either DNase 1 or corresponding vehicle. Each dot corresponds to a different thrombus fragment, n = 16 thrombi, each cut in half for paired comparison of both conditions.

In agreement with previous results^13^, we found that PAI-1, the main endogenous tPA inhibitor, was colocalized with NETs in IS thrombi (Figure 6A). Like plasminogen and tPA (Figures 2 and 3), α2AP, the primary inhibitor of plasmin, was found both in association with fibrin and NETs (Figure 6B). Quantification by ELISA of soluble PAI-1, protease nexin-1 (PN-1), neutrophil elastase (NE), and α2AP in supernatants recovered after the pre-incubation period of IS thrombi with either DNase 1 or vehicle (Figure 5A) showed that DNase 1 treatment had caused an increase in the release of PAI-1, PN-1, and NE from IS thrombi (Figure 6C-E). In contrast, α2AP levels were lower in the supernatants of DNase 1-treated thrombi (Figure 6F). Western blot analysis of α2AP integrity in reducing conditions revealed that, in addition to the band at the expected molecular weight for α2AP (70 kDa), IS thrombus supernatants also contained two α2AP fragments of approximately 39 and 30 kDa (Figure 6G). Remarkably, preincubation with DNase 1 led to a reduction in the higher 39 kDa fragment and to an increase in the smaller fragment, which migrated slightly lower than the initial 30 kDa band found in supernatants from vehicle-preincubated IS thrombi (Figure 6G). To verify whether these changes in α2AP integrity were associated with facilitation of fibrinolysis at the plasmin level, we assessed the impact of DNase 1 pretreatment on plasmin-induced IS thrombolysis. Pretreatment of IS thrombi with DNase 1 enhanced their lysis triggered by subsequent addition of plasmin (Figure 7A), showing that elimination of NETs facilitates fibrinolysis also at the plasmin level. In contrast, no benefits of DNase 1 pretreatment were observed when lysis was initiated by addition of streptokinase and plasminogen (Figure 7B), whose fibrinolytic activity is insensitive to tPA inhibitors and resistant to α2AP^24,25^.

**Figure 6.**
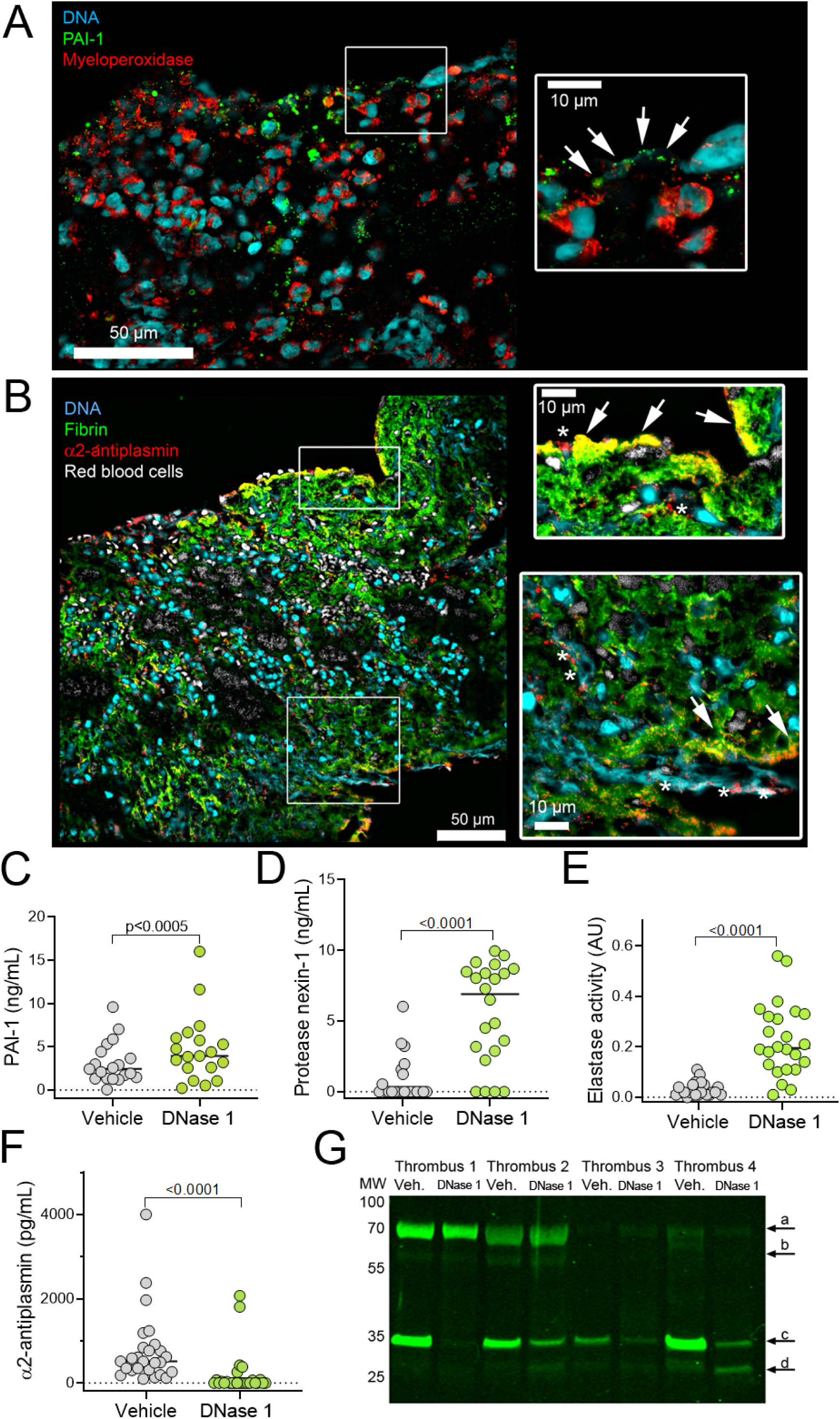
DNase 1 eliminates extracellular DNA-retained fibrinolysis inhibitors from acute ischemic stroke thrombi. **A.** Representative images of plasminogen activator inhibitor 1 (PAI-1, green) in ischemic stroke thrombi showing its association with strings of NETs (cyan, arrows, highlighted at higher magnification shown in the squared area) in a neutrophil-rich area at the thrombus periphery. Neutrophils were identified by the polynucleated aspect of their nuclei and positive staining for myeloperoxidase (red). **B.** Representative images of α2-antiplasmin (red) localization relative to fibrin (green, arrows) and NETs (cyan, asterisks). Higher magnification views of the squared areas are shown in the insets. **C-F.** Quantification of PAI-1, protease nexin-1, neutrophil elastase, and α2-antiplasmin, in ischemic stroke thrombus supernatants collected after 2 hours of incubation in HBSS-calcium-magnesium supplemented with either recombinant DNase 1 (pulmozyme, 100 µg/mL) or calcium saline (vehicle). Each dot represents the individual value of a different thrombus fragment. Each patient thrombus was split in 2 fragments, both fragments being randomly allocated to different treatment with either DNase 1 or vehicle. **G.** Western blot analysis (reducing conditions) of α2-antiplasmin integrity in supernatants from 4 different stroke thrombi cut in half and treated with either recombinant DNase 1 or calcium saline (vehicle). In addition to the expected 70 kDa α2-antiplasmin band (a), 3 additional lower molecular weight forms were detected (labeled b to d). Note that DNase 1 treatment was associated with a decrease in the 39 kDa form (c), which was mirrored by an increase in that of the lowest 30 kDa molecular weight band (d).

**Figure 7.**
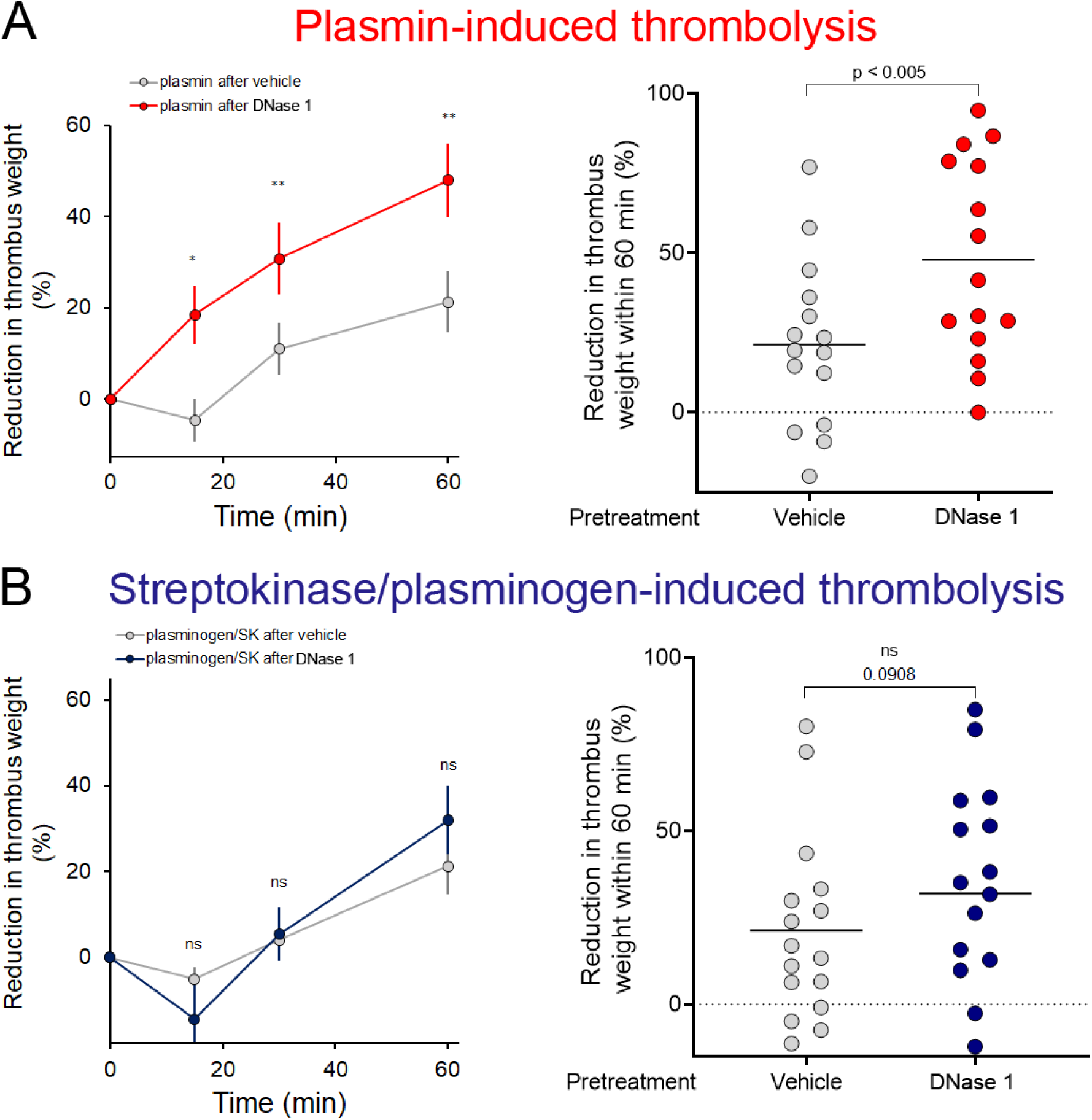
DNase 1 pretreatment facilitates subsequent thrombolysis of acute ischemic stroke thrombi by plasmin but not by plasminogen/streptokinase. A. Comparison of mean thrombus weight evolution (left panel) and of final reduction in thrombus weight (right panel) in response to plasmin added following a pretreatment of paired ischemic stroke thrombus fragments with either DNase 1 (pulmozyme, 100 µg/mL) or vehicle. n = 15 different thrombi, each cut in half, with each fragment being randomly allocated to either DNase 1 or vehicle pretreatment. * p > 0.05; ** p < 0.005; p values are for the comparison between the two groups at each time point. B. Comparison of mean thrombus weight evolution (left panel) and of final reduction in thrombus weight (right panel) in response to plasminogen/streptokinase added following a pretreatment of paired ischemic stroke thrombus fragments with either DNase 1 or vehicle. n = 14 different thrombi, each cut in half, with each fragment being randomly allocated to either DNAse 1 or vehicle pretreatment.

## Discussion

In the present study, we investigated the possible mechanisms underlying IVT failure in patients with AIS due to LVO. Seners *et al.* have shown recently that better collaterals were associated with early post-IVT recanalization (i.e. within 3 hours of IVT), whose incidence is particularly low in case of LVO AIS^1,5^. The association between better collaterals and higher recanalization efficacy of IVT could reflect improved delivery of tPA and plasminogen to the thrombus. It also suggests that failed IVT might result from inefficient tPA delivery. Nonetheless, previous histological studies comparing the features of AIS thrombi according to IVT status have reported consistent evidence that even when IVT fails to cause early recanalization, it is associated with signs of increased fibrinolysis, indicating at least partial tPA delivery. Signs of increased fibrinolysis in thrombi recovered from IVT patients notably included a reduced thrombus area and thinning of the thrombus fibrin network^26–29^. Our results add to the evidence that IVT-induced recanalization can fail despite tPA delivery and stimulation of fibrinolysis by directly showing that intrathrombus tPA and D-dimers content are increased in IVT patients compared to no-IVT patients. Taken together, while one cannot exclude collaterality-related delivery issues, these data suggest that failed IVT is due to insufficient (too-little, too-slow) rather than to absence of tPA delivery and fibrinolytic response.

In addition to tPA bound to well-defined scattered fibrin fibers in the thrombus core, we observed that tPA also accumulated in areas characterized by dense and compact fibrin, typical of previously described thrombolysis-resistant domains^10,30,31^. These regions with dense fibrin and tPA included the peripheral shell^10,32,33^, as well as neutrophil- and NETs-rich areas within the thrombus core. Using a microfluidic model of thrombosis comprising NETs and fibrin, we confirmed that tPA concentrated preferentially in NETs-rich areas. This suggests that NETs may act as decoy binding sites for tPA, concentrating it in fibrinolysis-resistant areas while titrating it away from « digestible » fibrin. In support of this hypothesis, we observed that tPA in AIS was involved in complexes including histones. It is worth noting that histones have been previously shown to interfere with fibrinolysis through non-covalent and covalent interactions with fibrin^34^. Our results add to growing evidence that histones might be critical negative regulators of fibrinolysis in AIS^12,34^.

Irrespective of IVT status, the vast majority of intrathrombus tPA was single chained and involved in several complexes, indicating that only a minor fraction of intrathrombus tPA might be freely available for promotion of plasminogen activation. Intriguingly, one of the complexes involving tPA also involved histone H1, supporting the possibility of direct interactions between NETs components and tPA. Nevertheless, the fact that addition of DNase 1 was sufficient to unlock fibrinolysis in the presence of plasminogen suggests that part of intrathrombus tPA is functional and not involved in irreversible inhibitory complexes, and can thus be mobilized for fibrinolysis. Furthermore, that DNase 1 converted the increased tPA content of IVT thrombi into increased fibrinolysis and thrombolyis bears clinical and pathophysiological implications. First, it suggests that DNase 1 could help improve the thrombolytic effect of current intravenously-administered therapeutic doses of alteplase and tenecteplase while, to date, the efficacy of DNase 1 as an adjuvant to tPA has been exclusively shown in experiments where tPA was delivered directly to AIS thrombi *ex vivo*. Moreover, it confirms our previous observations that NETs in AIS thrombi contribute to thrombolysis resistance not only by providing a secondary non-fibrin fibrillar scaffold to the thrombus but probably also by impairing tPA-dependent fibrinolysis^23^.

In their seminal study on the role of NETs in thrombosis, using an *in vitro* model of NETs-rich blood clots, Fuchs *et al.* showed that NETs could provide a secondary non-fibrin structural backbone to blood clots that compensated for the degradation of fibrin by tPA^35^. As a result, lysis of such NETs-rich blood clots required the combined use of tPA and DNase 1 for complementary degradation of both fibrin and DNA networks^35^. With respect to AIS, we have shown previously that within 1 hour of treatment, DNase 1 alone (i.e. without addition of exogenous tPA and plasminogen) had no thrombolytic effect towards AIS thrombi^9^. However, Peña-Martínez *et al.* showed that for longer extensive treatment periods (4 hours), DNase 1 alone exhibited a significant thrombolytic effect^14^. In line with the latter study, we show here that after 2 hours of treatment with DNase 1 alone, AIS thrombi had slightly but significantly lost weight. Yet, for 12 out of 14 AIS thrombi, this thrombolytic effect of DNase 1 was lost in the presence of aprotinin, thus demonstrating that, in most cases, it was largely dependent on plasmin and resulted from stimulation of the fibrinolytic potential of tPA and plasmin(ogen) present in AIS thrombi. Therefore, whereas DNase 1 likely exerts direct thrombolytic effects towards AIS thrombi with a particularly large NETs burden, our results indicate that it acts mainly indirectly by promoting fibrinolysis in AIS thrombi.

We then investigated the mechanisms underlying the indirect, fibrinolysis-dependent, thrombolytic effects of DNase 1. Locke and Longstaff have shown that, depending on their concentration, histones could stimulate plasminogen activation by tPA in solution^34^. To determine if fibrinolysis stimulation by DNase 1 relied on the release of factors capable of enhancing tPA and plasmin activity, we proceeded to sequential thrombolysis experiments in which fibrinolysis was triggered by addition of exogenous tPA, after DNase 1 treatment and washing of NETs degradation products and other potential factors released by DNase 1 from AIS thrombi. Our results show that potentiation of tPA-mediated thrombolysis by DNase 1 was maintained after washing of DNase 1 and soluble NETs degradation products. This argued against the involvement of profibrinolytic factors that would be released or generated by DNase 1. In contrast, it hinted at the possibility that DNase 1 lifted NETs-related antifibrinolytic activities. Several non-exclusive antifibrinolytic effects of NETs have been proposed previously. It has been shown that NETs-bound neutrophil elastase remains active^36^ and can degrade plasmin(ogen)^22^, limiting both plasmin formation and activity. Zhang *et al.* reported that circulating and intrathrombus NETs were decorated with PAI-1 in AIS patients^13^. In agreement with these results, we observed that PAI-1, as well as α2AP, were associated with NETs, and further showed that DNase 1 treatment led to the release of PAI-1, PN-1, and neutrophil elastase from AIS thrombi, thus indicating that NETs contribute to the retention of fibrinolysis inhibitors in AIS thrombi. Unlike tPA and plasmin(ogen), PAI-1, PN-1 and NE do not have specific binding domains for fibrin. Thus, upon NETs degradation by DNase 1, while PAI-1, PN-1 and NE are released from the thrombus, tPA and plasminogen might relocate onto fibrin where their interaction with fibrin protect them from inhibition. We further found that DNase 1 treatment caused the appearance of a α2AP fragment and enhanced thrombolysis by direct addition of plasmin, indicating that NETs in AIS thrombi block fibrinolysis both at the tPA and plasmin levels. That DNase 1 did not facilitate lysis by streptokinase/plasminogen stresses that the antifibrinolytic effects of NETs in AIS thrombi involve inhibitory pathways specific to tPA and plasmin. However, because in our experimental settings, AIS thrombi were cut in half, giving drugs access to the thrombus core, one cannot exclude the involvement of non-specific antifibrinolytic effects of NETs, such as impairment of drug diffusion through densification of the fibrin network^22,34^. In conclusion, our results indicate that although IVT succeeds in increasing tPA in AIS thrombi, NETs limit its fibrinolytic efficacy by decreasing tPA and plasminogen access to fibrin and by promoting intrathrombus retention of fibrinolysis inhibitors.

Finally, by showing that DNase 1 and tPA can act sequentially, with DNase 1 pretreatment facilitating the subsequent action of tPA, our results provide valuable information for refining and optimizing the design of clinical trials aimed at assessing the efficacy of DNase 1 as an adjuvant to tPA. DNase 1 administered within 24 hours of stroke onset has proven particularly safe, with no hemorrhagic complications in animal model of ischemic stroke^37,38^, and even antihemorrhagic and beneficial effects in tPA-treated mice^39,40^ and models of subarrachnoid hemorrhage^41,42^. The safety of acute administration of DNase 1 in models of ischemic and hemorrhagic stroke combined with its ability to waive NETs-related antifibrinolytic effects and clear the way for tPA suggest that it could possibly be administered very early, i.e. before imaging, to stimulate both endogenous and IVT-induced fibrinolysis.

## Acknowledgements

This work was realized with the help of grants from the ANR (# ANR-18-RHUS-0001, RHU Booster, ANR-22-CE17-0032 INFLAME, ANR-23-CE17-0023-01 DELIASE), Fondation de France (FDF 00086496), and Fondation pour la Recherche Médicale (#DPC20171138959), and INSERM (MESSIDORE SAVE-BRAIN).

